# Leveraging AI to Explore Structural Contexts of Post-Translational Modifications in Drug Binding

**DOI:** 10.1101/2025.01.14.633078

**Authors:** Kirill E. Medvedev, R. Dustin Schaeffer, Nick V. Grishin

## Abstract

Post-translational modifications (PTMs) play a crucial role in allowing cells to expand the functionality of their proteins and adaptively regulate their signaling pathways. Defects in PTMs have been linked to numerous developmental disorders and human diseases, including cancer, diabetes, heart, neurodegenerative and metabolic diseases. PTMs are important targets in drug discovery, as they can significantly influence various aspects of drug interactions including binding affinity. The structural consequences of PTMs, such as phosphorylation-induced conformational changes or their effects on ligand binding affinity, have historically been challenging to study on a large scale, primarily due to reliance on experimental methods. Recent advancements in computational power and artificial intelligence, particularly in deep learning algorithms and protein structure prediction tools like AlphaFold3, have opened new possibilities for exploring the structural context of interactions between PTMs and drugs. These AI-driven methods enable accurate modeling of protein structures including prediction of PTM-modified regions and simulation of ligand-binding dynamics on a large scale. In this work, we identified small molecule binding-associated PTMs that can influence drug binding across all human proteins listed as small molecule targets in the DrugDomain database, which we developed recently. 6,131 identified PTMs were mapped to structural domains from Evolutionary Classification of Protein Domains (ECOD) database. **Scientific contribution.** Using recent AI-based approaches for protein structure prediction (AlphaFold3, RoseTTAFold All-Atom, Chai-1), we generated 14,178 models of PTM-modified human proteins with docked ligands. Our results demonstrate that these methods can predict PTM effects on small molecule binding, but precise evaluation of their accuracy requires a much larger benchmarking set. We also found that phosphorylation of NADPH-Cytochrome P450 Reductase, observed in cervical and lung cancer, causes significant structural disruption in the binding pocket, potentially impairing protein function. All data and generated models are available from DrugDomain database v1.1 (http://prodata.swmed.edu/DrugDomain/) and GitHub (https://github.com/kirmedvedev/DrugDomain). This resource is the first to our knowledge in offering structural context for small molecule binding-associated PTMs on a large scale.

## Introduction

Post-translational modifications (PTMs) play a crucial role in regulating protein activity, stability, and function. PTMs can significantly influence a protein’s interactions and overall functional activity by introducing new chemical functionalities and altering their structural and electrostatic properties. While the majority of PTMs indeed modulate protein interactions, some, such as certain types of glycosylation, may primarily affect protein stability, folding, or trafficking without directly influencing binding partners [1, 2]. By modulating these properties, PTMs contribute to cellular signaling, metabolic pathways, and the dynamic response of proteins to environmental and physiological changes [3, 4]. PTMs provide a level of functional diversity that surpasses the inherent properties of the 20 standard amino acids. They introduce a wide array of chemical groups, including phosphates, sugars, lipids, and small molecules, expanding the chemical repertoire of proteins. This expanded chemical repertoire enables, for example, new binding specificities, as phosphorylation can create novel sites for protein-protein interactions [5, 6]. PTMs can also direct proteins to specific cellular compartments. For example, palmitoylation adds lipid groups to proteins, facilitating their membrane association [7]. The evolutionary advantage of PTMs lies in their ability to rapidly and reversibly modulate protein function in response to changing cellular conditions. This dynamic regulation allows organisms to adapt to environmental challenges, respond to signals, and fine-tune cellular processes with precision [8, 9]. Therefore, PTMs play a crucial role in the development and progression of various diseases, including cancer, neurodegenerative disorders, and diabetes [10, 11]. Recent advances in mass spectrometry have revolutionized the study of PTMs, enabling the identification and characterization of hundreds of distinct PTM classes across entire proteomes [12, 13]. However, assessing the functional relevance of each PTM remains a significant challenge.

In recent years, the development of accurate AI-based methods for predicting the structure of complex protein systems has revolutionized computational structural biology [14]. These prediction methods allow for the exploration of the structural context of PTMs on a proteome-wide scale, which was previously impossible [15, 16]. Various resources provide structure-related information about PTMs, including StructureMap [15] and Scop3P [17]. PTMs can significantly impact the affinity of drug binding by altering the protein’s structure and electrostatic properties. For example, phosphorylation can introduce new charge groups, affecting electrostatic interactions between the protein and the drug, and may induce conformational changes in the protein [18, 19]. Glycosylation can affect a drug’s binding affinity to receptors by altering the structure of the glycans on the drug [20–22]. A diverse array of PTMs, including ubiquitination, hydroxylation, methylation, acetylation, and phosphorylation, serve as critical regulatory mechanisms for undruggable transcription factors, modulating their stability, subcellular localization, protein-protein interactions, and DNA-binding specificity [23, 24]. Given the challenges associated with directly targeting undruggable transcription factors, modulating their activity through PTM-based approaches presents a viable alternative [25]. For example, inhibiting JAKs (Janus kinase) provides an effective therapeutic strategy for diseases driven by aberrant JAK/STAT signaling, as JAKs directly phosphorylate and activate STAT proteins, crucial for pathway activation [26, 27]. On a large scale, the potential impact of PTMs located within proximity of the binding site on drug-binding affinity has been predicted by several resources, including, but not limited to, dbPTM [28], canSAR [29], CruxPTM [30]. However, the structural aspects of these PTMs on small molecule binding have not been extensively studied. Here, we address this gap using state-of-the-art AI-based methods.

Specifically, we focused on the DrugDomain database, which we recently developed, containing interactions between human protein domains and small molecules. This database includes both experimentally determined PDB structures and AlphaFold models enriched with ligands from experimental data [31]. For these human proteins, we identified small molecule binding-associated PTMs that occurred within 10 Å of the small molecule and generated models of the modified structures using AlphaFold3 [32], RoseTTAFold All-Atom (RFAA) [33], Chai-1 (v0.1.0) [34], and KarmaDock [35]. We mapped identified PTMs to structural domains from the Evolutionary Classification of Protein Domains (ECOD) database [36, 37], providing valuable data for exploring evolutionary aspect of PTMs [9]. Our structural models revealed that phosphorylation of NADPH-Cytochrome P450 Reductase, which was detected in cervical and lung cancer, causes significant structural disruption in the binding pocket and potential dysfunction of this protein. We have reported these data on GitHub (https://github.com/kirmedvedev/DrugDomain) and on the DrugDomain database v1.1 (http://prodata.swmed.edu/DrugDomain/), which is the first resource to provide structural context of small molecule binding-associated PTMs on a large scale.

## Materials and Methods

### Identification of small molecule binding-associated PTMs

Post-translational modifications (PTMs) were retrieved from the dbPTM database (August 2024 version), which integrates more than 40 smaller PTM-related databases and reports more than 2 million experimental PTM sites [28]. Affinity for small molecule binding can be affected by PTMs occurring within 10 Å of the small molecule [38, 39]. Therefore, using BioPython [40] we identified PTMs within 10 Å of all atoms of each small molecule bound to human proteins in the DrugDomain database [31]. The DrugDomain database documents interactions between protein domains and small molecules both for experimentally determined PDB structures and AlphaFold models which were modelled with ligands from experimental structures based on protein sequence and structure similarity using AlphaFill approach, which transplants missing small molecules and ions into predicted AlphaFold models based on sequence and structure similarity [41]. In cases where small molecule binding-associated PTMs were identified in a PDB structure, we generated a BLAST [42] alignment for the sequence of the PDB chain containing the PTMs against the UniProt sequence to determine the UniProt numbering of the residues with the PTM. Chimeric PDB structures where a PDB chain includes multiple UniProt accessions were excluded. We observed cases where the number and type of PTM-containing residue in the dbPTM database did not match UniProt sequence and numbering. We disregarded these cases and excluded them from further analysis. The overall non-duplicated number of identified small molecule binding-associated PTMs is 6,131. The non-duplicated number of PTMs is determined by counting PTMs per UniProt accession. Counting PTMs per ECOD domain introduces duplications, as multiple PDB structures often correspond to the same UniProt accession. We analyzed each protein-ligand pair and documented all PTMs occurring within 10 Å of the ligand in the DrugDomain database. 6,131 includes 30 types of PTMs (such as phosphorylation, ubiquitination, etc.) and 47 combinations of PTM and amino acid types (for example Phosphorylation of SER, Acetylation of LYS, etc.) (Supplementary Table 1). Identified small molecule binding-associated PTMs were mapped to structural domains from ECOD database v292 [36, 37].

### Modelling of protein structures with small molecule binding-associated PTMs

Overall, we utilized four approaches to create protein models with PTMs and small molecules: AlphaFold3 server [32], RoseTTAFold All-Atom (RFAA) [33], Chai-1 (v0.1.0) [34] and KarmaDock [35]. To test the selected methods, we targeted proteins where phosphorylation sites within 12 Å of the small molecule-binding site are likely to influence small molecule binding affinity [18]. For this test set, we created models of 64 combinations of protein targets and drugs with PTMs and 60 combinations of unmodified protein targets and drugs (several proteins in this set contain two PTMs), using RFAA, Chai-1, KarmaDock, and AlphaFold3 (Supplementary Table 2). Each modeling run for the test set was performed three times, except AlphaFold3 (one time). Different methods produce varying numbers of output models per one run: RFAA – one model per run, Chai-1 – five, KarmaDock – three, AlphaFold3 - five. Thus, the total number of modeled structures retained per unmodified protein-ligand pair in the test set is: RFAA – 3, Chai- 1 – 15, KarmaDock – 9, and AlphaFold3 - 5 with the same numbers applied to the PTM-modified state. For our dataset of identified small molecule binding-associated PTMs we used AlphaFold3, RFAA and Chai-1 for creating models. KarmaDock was used only in the test set and for the examples discussed in this manuscript. We used protein models containing the PTM generated by Chai-1 for KarmaDock input. Each modeling run for our dataset was performed once, except for the examples discussed in this manuscript (which were run three times). Thus, the total number of modeled structures retained per protein-ligand pair in our dataset is: AlphaFold3 – 5, RFAA – 1, Chai-1 – 5, KarmaDock – 3. These generated protein models will be compared to available experimental structures, as described in the next subsection. For AlphaFold3 server runs, we used the complete protein sequence from UniProt KB [43]. For RFAA and Chai-1 runs, in cases where proteins exceeded 1,500 amino acids, we used the PDB chain sequence or the sequence of the ECOD domain interacting with the small molecule. RFAA runs require small molecules and the chemical group attached as a PTMs to be provided as SDF files. All required SDF files were obtained from RCSB Protein Data Bank [44]. SDF files of the chemical group attached as PTMs were manually modified where necessary to handle “leaving groups”, as recommended by the RFAA manual. Chai-1 runs require SMILES formulas of small molecules, which were retrieved from DrugBank [45], and CCD codes of modified residue, which were obtained from the Chemical Component Dictionary [46]. For all modelling runs randomly assigned seeds were used. We additionally tested DiffDock [47] and FeatureDock [48] docking methods, however we found that current versions of these methods cannot process PTMs in the protein structures. Due to technical limitations of each selected method, we created protein models with PTMs and small molecules for 27 combinations of amino acid and small molecule binding-associated PTM types (Supplementary Table 3). Overall, we obtained 1,041 AlphaFold3 models, 9,169 RFAA models and 3,968 Chai-1 models. All models can be accessed through DrugDomain database website (http://prodata.swmed.edu/DrugDomain/).

### Calculation of Root Mean Square Deviation (RMSD)

To evaluate the potential effect of PTMs on the binding mode of small molecules, we calculated the RMSD between the modeled position of the molecule and its experimentally determined position or the position predicted by AlphaFill (see above). Calculation of RMSD was conducted using PyMOL [49]. First, modeled and PDB/AlphaFill structures were aligned using PyMOL “align” function, which takes into account sequence similarity. The align function begins by performing a global dynamic-programming sequence alignment on a per-residue basis for the input atom selections, utilizing the BLOSUM62 scoring matrix from BLAST. Next, it establishes a correspondence between atoms in the selections, including matching side-chain atoms if specified in the selection arguments. An initial superposition is conducted, followed by up to five cycles of iterative refinement. During each cycle, atoms with deviations exceeding two standard deviations from the mean are excluded, and the fitting process is repeated. In cases where the orientation of domains in a multidomain protein model does not match the domain orientation in the experimental structure (Supplementary Fig. 1), only the domains involved in small molecule binding were used for structural alignment. After the alignment of structures PyMOL “rms_cur” function was used to calculate RMS difference for atoms of modeled and PDB/AlphaFill small molecule. Rms_cur computes the RMS difference between two atom selections without performing any fitting. If PDB/AlphaFill structure contain more than one small molecule of interest, RMSD calculations were conducted for each molecule. This approach of RMSD calculation requires matching atom names between modeled and PDB/AlphaFill structures. RFAA’s and KarmaDock’s output models contain small molecule atom names that do not match the atom names in the original CIF files; however, the order of these atoms remains the same. Thus, before RMSD calculation, small molecule atoms in the RFAA and KarmaDock models were renamed according to the CIF small molecule files obtained from the RCSB PDB. Chai-1 output models contain small molecule atom names and order that do not match the atom names and order in the CIF files. We used manual approach to map atom names in Chai-1 models to atom names in CIF files. Thus, we calculated RMSD of Chai-1 models only for test set. Scripts for RMSD calculations are available at GitHub (https://github.com/kirmedvedev/DrugDomain).

### Calculation of Local Distance Difference Test for Protein–Ligand Interactions (lDDT-PLI)

Additionally we calculated lDDT-PLI score that assesses the conservation of contacts between the ligand and the protein, comparing experimental structure and PTM-modified model [50]. First, we identified the interface atoms by selecting protein and ligand atoms that lie within 5 Å. from any atom of the binding partner. For each interface atom in the reference structure, we calculated the distances to its neighboring interface atoms (which may include both protein and ligand atoms) and did the same for the corresponding atoms in predicted PTM-modified model. For each pair of interface atoms *i* and *j*, we computed the absolute difference between the distance in the reference structure, 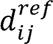, and the corresponding distance in predicted model, 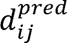. For each pair of atoms, we used a threshold based scoring function *f* that assigns a value between 0 and 1 based on how close the two distances are:

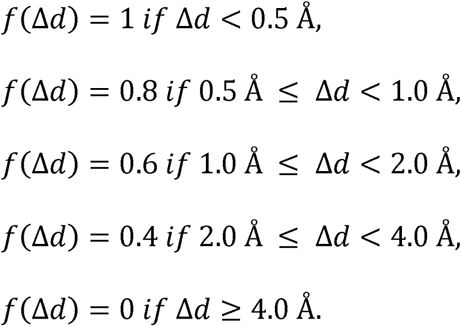

For each interface atom, we averaged the *f* values over all its considered pairs. Then, the overall lDDT□PLI score is the average over all interface atoms:

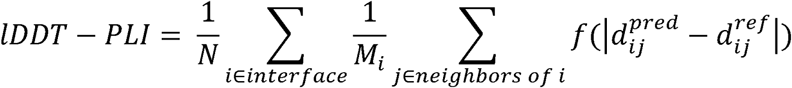

Where *N* is the number of interface atoms and *M_i_* is the number of pairs considered for atom *i*.

## Results and Discussion

### Distribution of small molecule binding-associated PTMs in ECOD domains

We defined small molecule binding-associated PTMs as those located within 10 Å of a small molecule (see Materials and Methods). To identify these PTMs, we utilized the dbPTM database [28] and analyzed all human proteins we previously reported in the DrugDomain database [31]. The total number of unique small molecule binding-associated PTMs identified is 6,131. This comprises 30 types of PTMs (e.g., phosphorylation, ubiquitination) and 47 specific combinations of PTM types and amino acid residues (e.g., phosphorylation of serine, acetylation of lysine) (Supplementary Table 1). We mapped identified PTMs to structural domains from the ECOD database [36, 37]. Figure 1 shows the distribution of small molecule binding-associated PTMs in protein domains at the highest level of ECOD classification – architecture groups (A-groups). In ECOD, we utilize 21 architecture (A-group) levels to provide a broad classification system for domains, focusing on their secondary structure content, overall structural arrangement, and potential functional roles. In the DrugDomain database, we document interactions between human protein domains and small molecules not only for experimentally determined PDB structures but also for AlphaFold models enriched with ligands from experimental structures. This enrichment is achieved using the AlphaFill approach [41], which transplants missing small molecules and ions into predicted protein models based on sequence and structure similarity.

**Figure 1.**
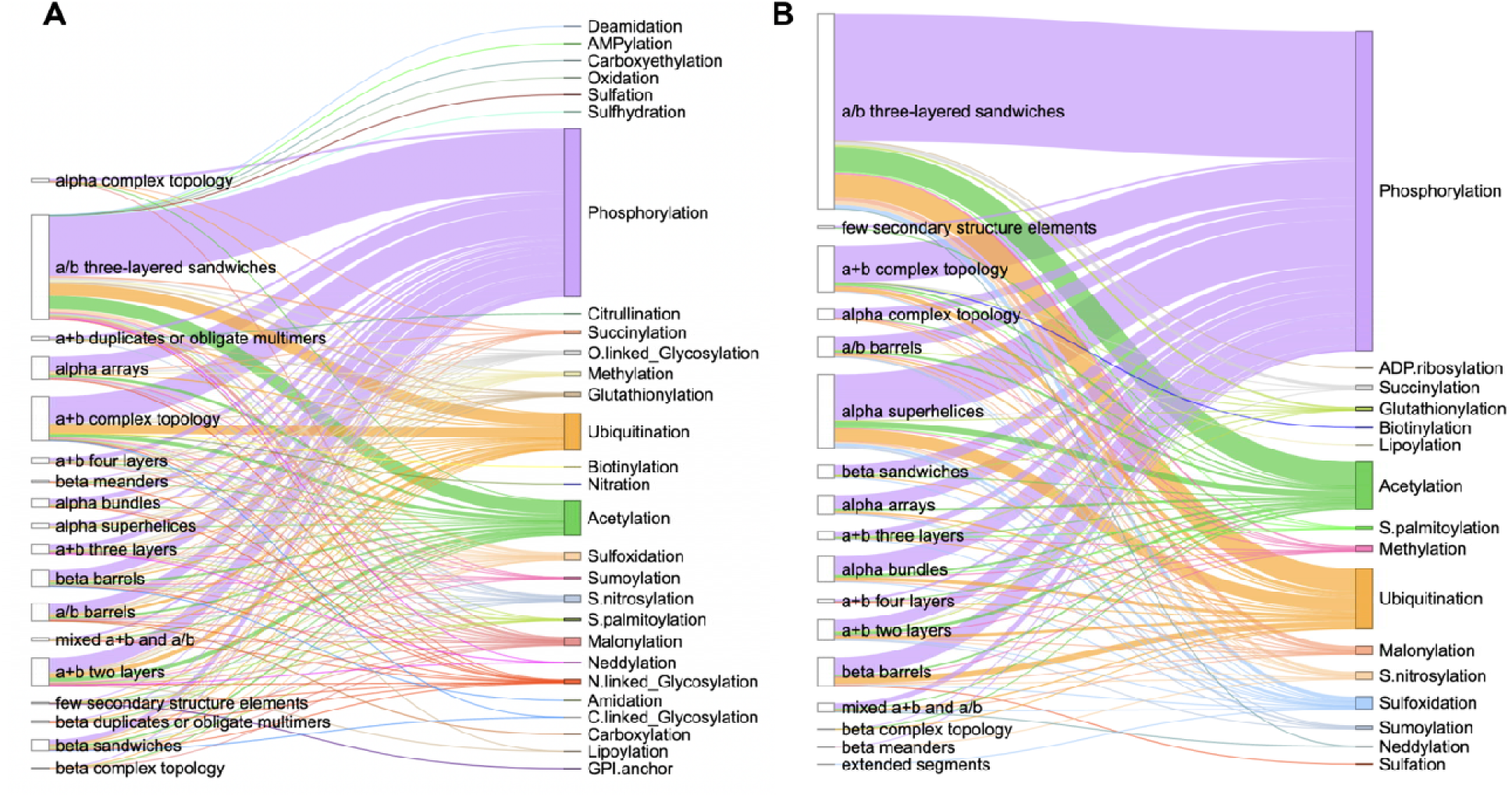
Distribution of small molecule binding-associated PTMs types in ECOD architecture groups. **(A)** Statistics for experimental PDB structures. **(B)** Statistics for AlphaFill models. The length of each vertical line represents the number of PTMs per ECOD A-group.

Thus, Figure 1 shows separate statistics for experimental PDB structures (Fig. 1A) and for AlphaFill models (Fig. 1B). As expected, the top three most prevalent types of small molecule binding-associated PTMs are phosphorylation, ubiquitination, and acetylation. Phosphorylation is considered the most prevalent type of PTM due to its highly reversible nature, which allows for rapid and dynamic regulation of protein function, making it ideal for cellular signaling pathways that need to quickly respond to changing stimuli; it can easily activate or deactivate proteins by adding a phosphate group, impacting various cellular processes like cell growth, differentiation, and apoptosis [51]. Ubiquitination is a highly versatile regulatory mechanism, allowing cells to control a wide range of cellular processes by targeting proteins for degradation, altering their activity and acting as a key switch for various biological pathways [52]. Finally, acetylation plays a central role in regulating fundamental biological processes. It is critical in gene expression through the acetylation of histone proteins, influences protein function by modulating their activity and regulates cellular metabolism [53].

The top three ECOD A-groups with the largest number of small molecule binding-associated PTMs across experimental PDB structures include a/b three-layered sandwiches, a+b complex topology, and a+b two layers (Fig. 1A). Proteins that comprise the majority of a/b three-layered sandwiches architecture adopt a Rossmann-like fold. We previously demonstrated that these proteins are among the most ubiquitous structural units in nature and are key elements in many metabolic pathways [54, 55]. The architecture group a+b complex topology encompasses various types of protein kinases, which play a critical role in cellular signaling by phosphorylating other proteins. These kinases are not only central to regulating diverse biological processes, such as cell division, metabolism, and apoptosis, but they are also subject to multiple PTMs themselves. These PTMs, including phosphorylation, acetylation, and ubiquitination, modulate kinase activity, stability, and interaction networks, further enhancing their functional versatility and regulatory capacity [56, 57]. Finally, a+b two layers architecture includes heat shock proteins (HSP) which play a critical role as molecular chaperones. PTMs can directly modulate the chaperone activity of HSPs, either enhancing or inhibiting their ability to bind and refold unfolded proteins [58]. This A-group also includes SH2 domains of proto-oncogene tyrosine-protein kinase Src, where specific PTM sites function as critical regulatory elements. For example, phosphorylation at key tyrosine residues within these sites serves as an inhibitory mechanism, maintaining Src in an inactive state by stabilizing intramolecular interactions that suppress its kinase activity [59].

The number of small molecule binding-associated PTMs types obtained from PDB structures (Fig. 1A) is greater than that from AlphaFill models (Fig. 1B). This may be explained by the fact that the AlphaFill approach derives ligands from the experimental structures from Protein Data Bank. However, one small molecule binding-associated PTM type is present among AlphaFill models and absent among PDB set – ADP-ribosylation (Fig. 1B). ADP-ribosylation is a reversible process that involves adding ADP-ribose units to a protein, that regulates various cellular functions [60]. In our dataset, ADP-ribosylation of cysteine located within 10 Å of the ligand was identified in two mitochondrial proteins - Glutamate dehydrogenase 1 (P00367) and 2 (P49448). ADP-ribosylation of CYS172 has been reported in both proteins; however, the functional relevance of these PTMs remains unclear [61].

### Chai-1 and RoseTTAFold All-Atom demonstrate the ability to predict the effects of PTMs on small molecule binding

To evaluate approaches for modeling protein structures with PTMs and their potential impact on small molecule binding, we analyzed protein targets where phosphorylation sites within 12 Å of the small molecule-binding site are likely to affect binding affinity [18]. While this list does not represent ground truth, it includes cases where PTM sites are highly likely to influence the protein’s function and binding affinity. For this test set, we generated models for 64 distinct combinations of protein targets and drugs, incorporating both PTM-modified and unmodified states (Supplementary Table 2). The modeling and docking were conducted using RoseTTAFold All-Atom (RFAA) [33], Chai-1 [34] and KarmaDock [35]. To evaluate the performance of the selected methods we calculated Root Mean Square Deviation (RMSD) of the ligands between modeled and experimental structure after the alignment of protein structures. To calculate RMSD we compared unmodified (and PTM-modified where available) experimental structure with PTM-modified and unmodified models. The ligand RMSD values were averaged for each case (with each modeling run performed three times) and compared between the PTM-modified and unmodified states. The authors of the test list identified two classes of phosphorylation site effects: Class 1, where phosphorylation inhibits both drug binding and target activity, and Class 2, where phosphorylation may reduce drug affinity without significantly inhibiting target function, and in some cases, may actually increase activity [18]. Thus, one would expect the ligand RMSD of the PTM-modified state to be higher than that of the unmodified state, at least for Class 1 cases.

Our results revealed that models generated by RFAA and Chai-1 predicted ligand positions in unmodified states that were close to the experimental positions. Moreover, for 13% of cases (8 out of 64) these methods predict higher ligand RMSD for PTM-modified states (Fig. 2A, B). However, in several cases, Chai-1 models exhibited a high standard deviation, indicating inconsistency in predictions for both unmodified and PTM-modified states (Fig. 2B). KarmaDock did not demonstrate high accuracy in predicting ligand positions in unmodified states for the studied test set (Fig. 2C). AlphaFold3 demonstrated high accuracy in predicting ligand positions in unmodified states; however, in most cases, ligand positions remained unchanged after introducing PTMs (Fig. 2D). The example of the case when RFAA and Chai-1 both predicted higher ligand RMSD for PTM-modified states is shown in Figure 3.

**Figure 2.**
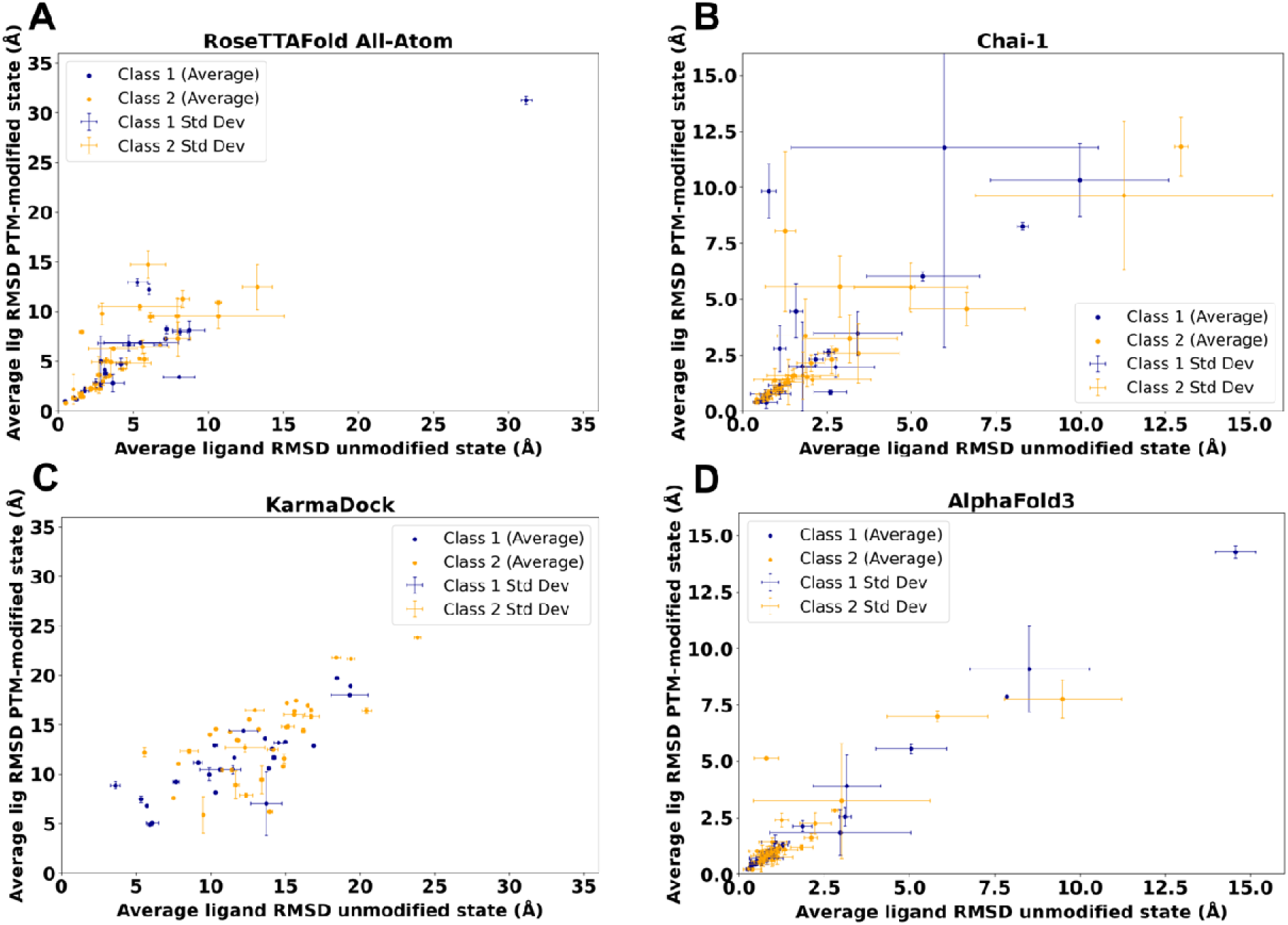
Average ligand RMSD for the PTM-modified and unmodified states in model generated by different approaches. **(A)** RoseTTAFold All-Atom (RFAA). **(B)** Chai-1. **(C)** KarmaDock. **(D)** AlphaFold3.

**Figure 3.**
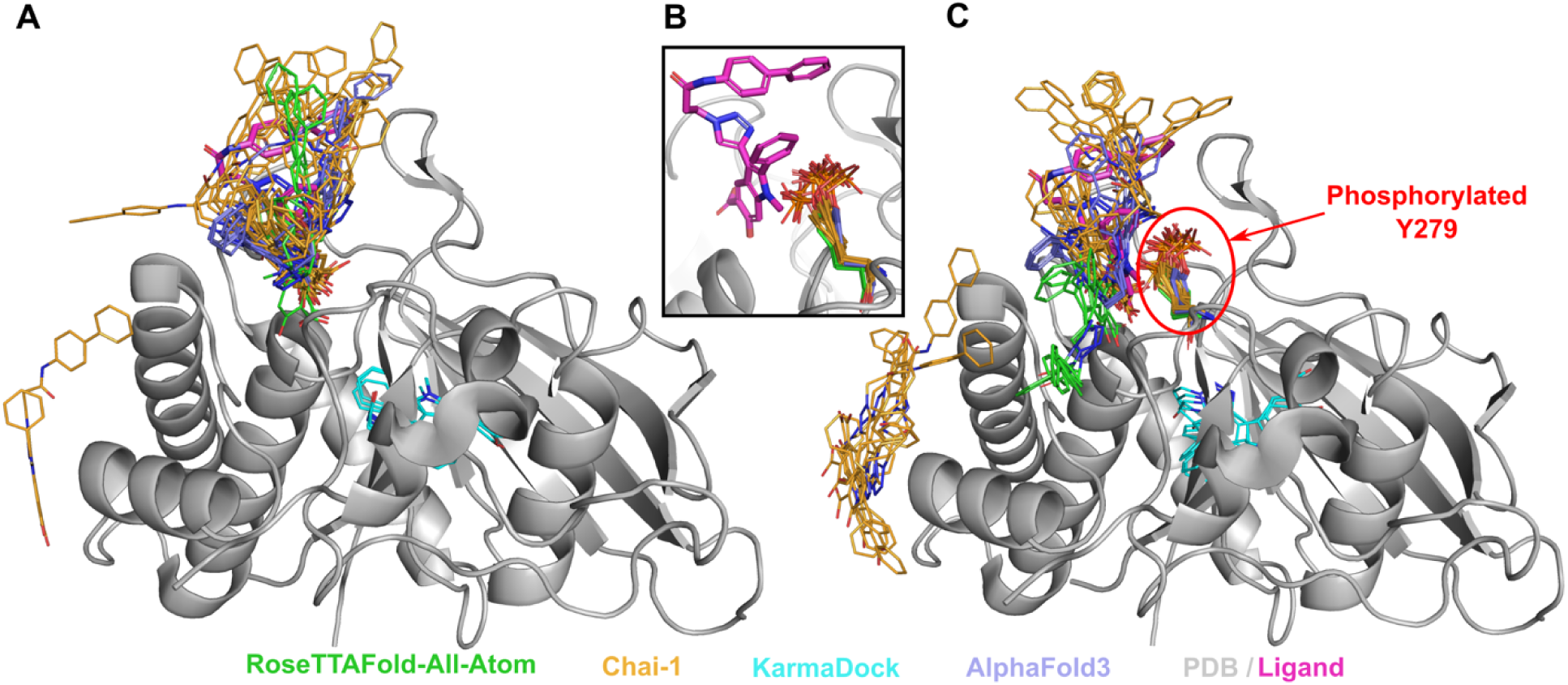
Structure of Tyrosine-protein phosphatase non-receptor type 11 (SHP-2) (PDB: 3O5X, shown in grey) in complex with inhibitor (II-B08, in magenta) and the modelled positions of this drug. **(A)** Unmodified state. **(B**) Zoomed-in view of modelled PTMs and experimental drug position. **(C)** PTM-modified state. Drug positions modelled by RFAA shown in green, Chai-1 in orange, KarmaDock in cyan, AlphaFold3 in slate. Experimental position of the drug is shown in magenta and thick sticks. The phosphorylated residue is shown in colors corresponding to the methods by which it was modeled.

Tyrosine-protein phosphatase non-receptor type 11 (SHP-2) (PDB: 3O5X, UniProt: Q06124) plays an important role in growth factor and cytokine signaling [62]. It was shown that inhibitor B08 (compound 9) conducts chemical inhibition of SHP-2 that may be therapeutically useful for anticancer and antileukemia treatment [63]. AlphaFold3, RFAA and Chai-1 approache predicted positions of the drug similar to the experimental (except one run of Chai-1) (Fig. 3A). Phosphorylation of SHP-2 on Y279 that is important for keeping SHP-2 in an inactive state [64]. The introduction of this PTM, located very close to the binding site, into structural models showed significant differences in the drug positions predicted by RFAA and Chai-1– most predicted molecules are located outside of the binding pocket (Fig. 3B). However, in the PTM-modified state, the ligand’s position modeled by AlphaFold3 remained unchanged. This case belongs to Class 1 phosphorylation site effects. Another example of Class 1 phosphorylation site effects is shown in Figure 4. However, in this case none of the methods predicted change of the drug’s binding mode.

**Figure 4.**
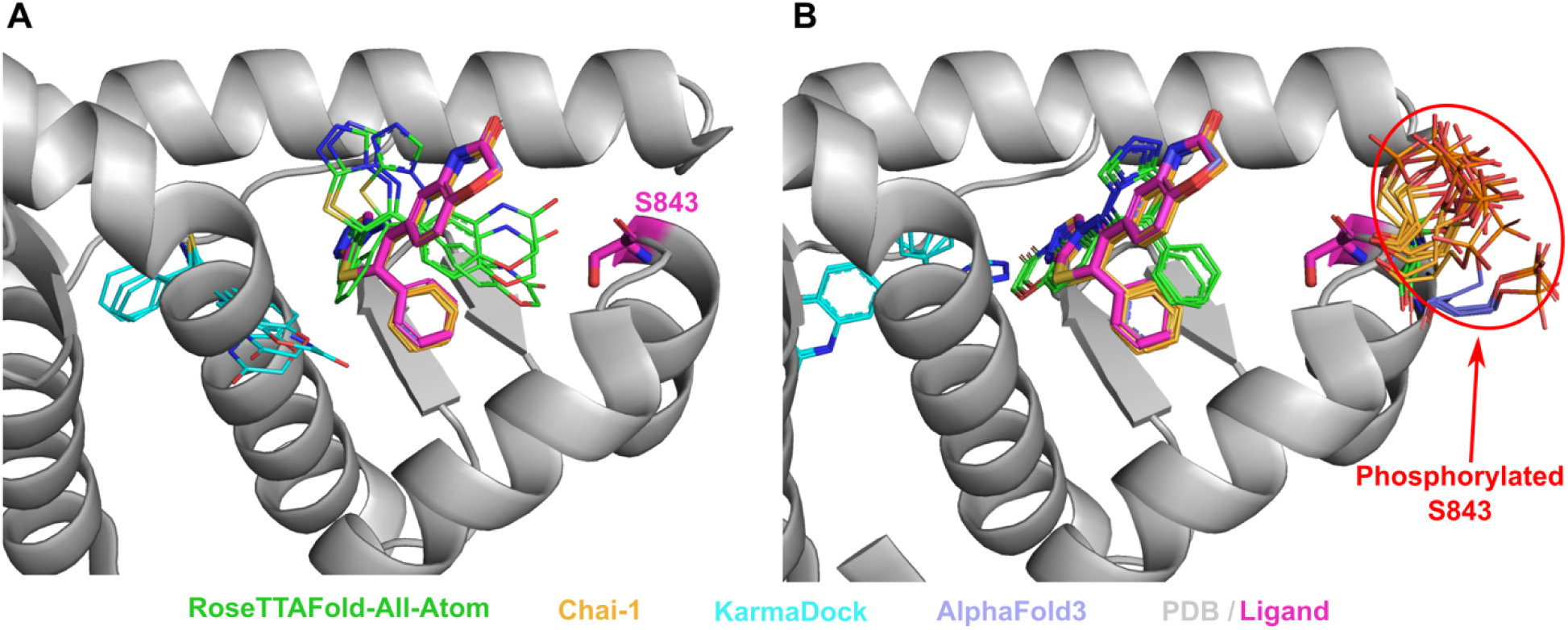
Structure of Mineralocorticoid receptor (PDB: 3VHV, shown in grey) in complex with inhibitor (PDB id: LD1, in magenta) and the modelled positions of this drug. **(A)** Unmodified state. **(B)** PTM-modified state. Drug positions modelled by RFAA shown in green, Chai-1 in orange, KarmaDock in cyan, AlphaFold3 in slate. Experimental position of the drug and Ser843 are shown in magenta and thick sticks. The phosphorylated residue is shown in colors corresponding to the methods by which it was modeled.

It was discovered that phosphorylation of mineralocorticoid receptor at Ser843 reduces the affinity for the natural agonist and inactivates the receptor [65]. Phosphorylation at the binding site for both the agonist and inhibitor of the mineralocorticoid receptor suggests that phosphorylation of Ser843 likely reduces drug affinity [18]. However, modeled structures did not reveal any difference in drug’s binding mode between PTM-modified and unmodified states (Fig. 4A, B). The modeled phosphorylated serine residues point outside the binding pocket (Fig. 4B), whereas it has been suggested that they should point toward the pocket, thereby preventing inhibitor binding and deactivating the receptor [18].

Most kinases from the test list fall into Class 2 category when phosphorylation inhibits drug binding while activating or not significantly inhibiting the target function [18]. Inactive state of insulin-like growth factor 1 receptor can bind inhibitor when the activation loop (Fig. 5A and 5B, shown in salmon) is located close to the binding site. Phosphorylation of Tyr1161 significantly reduces the affinity of inhibitor binding, which causes the activation and brings activation loop (Fig. 5A and 5B, shown in green) far from the binding site [66–68]. Our modeling results revealed that RFAA and Chai-1 do not differentiate between the active and inactive states of this protein, whereas AlphaFold3 does. All modeling runs of RFAA and Chai-1, for both unmodified and PTM-modified states, resulted in the activation loop being positioned in a manner corresponding to the active state (Fig. 5A, B), and only AlphaFold3 captured inactive state correctly (Fig. 5A – shown in light blue). The position of phosphorylated Tyr1161 modeled by AlphaFold3 and Chai-1 was closer to the experimentally observed modification in the insulin receptor (PDB: 1IR3) than the position modeled by RFAA (Fig. 5B). However, no significant differences were observed in the drug binding mode between the models of unmodified and PTM-modified states. Finally, the phosphorylation of Ser222 in mitogen-activated protein kinase 1 (MAP2K1) is an important mechanism for regulating its activity [69, 70]. Our models (RFAA and Chai-1) indicated a significant change in the binding mode of the MAP2K1 inhibitor [71] in the PTM-modified state (Fig. 5C, D), consistent with previous suggestions [18], whereas ligand’s position modeled by AlphaFold3 remained unchanged.

**Figure 5.**
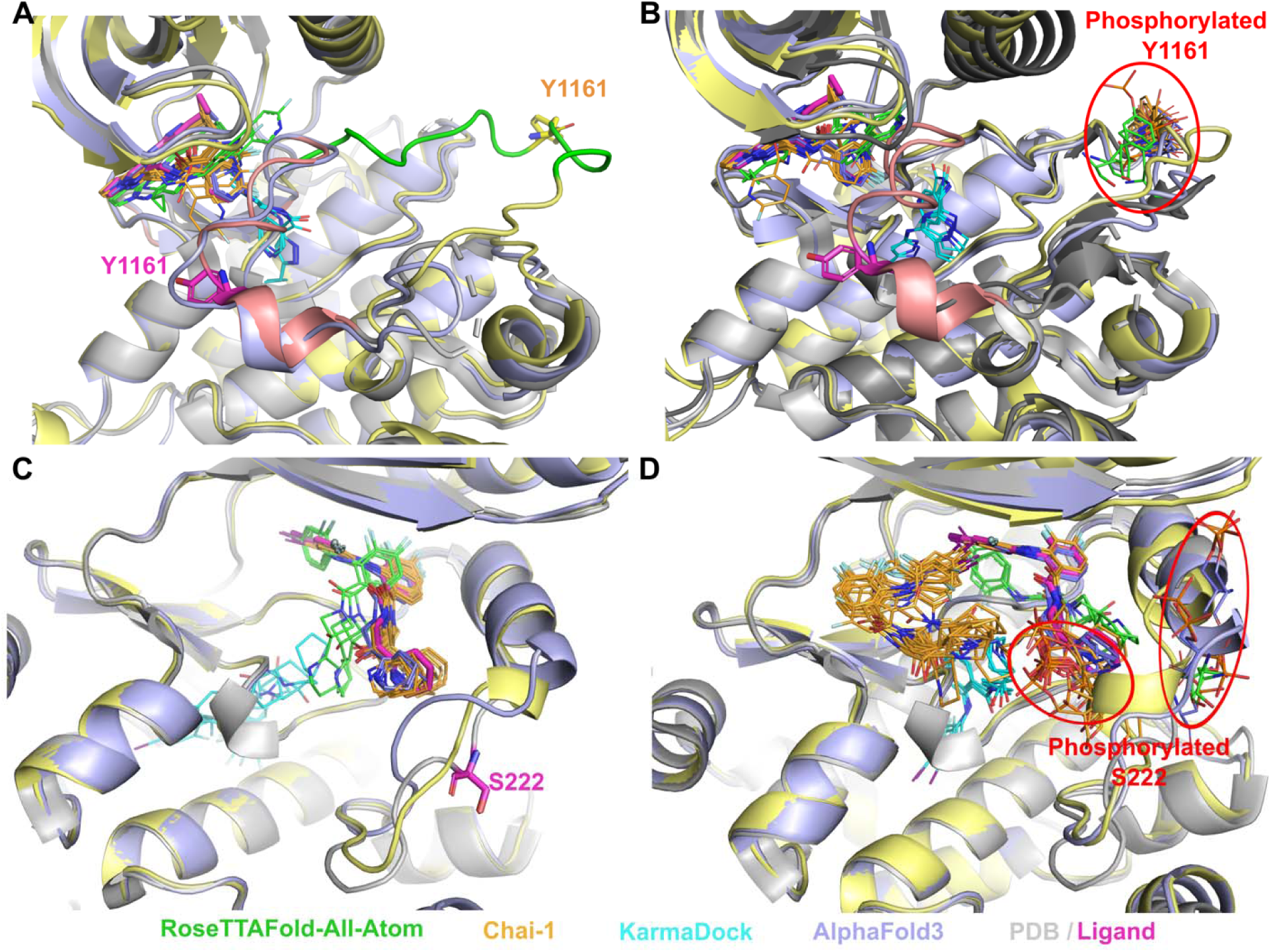
Examples of Class 2 phosphorylation site effects. **(A)** Unmodified state of inactive insulin-like growth factor 1 receptor (PDB: 3NW7, shown in grey). **(B)** PTM-modified state of inactive insulin-like growth factor 1 receptor. Phosphorylated insulin receptor (PDB: 1IR3) shown in dark grey. Experimental position of the drug (PDB id: LGV) and Tyr1161 are shown in magenta and thick sticks. Activation loop of inactive receptor is shown in salmon, active and modeled in green. **(C)** Unmodified state of mitogen-activated protein kinase 1 (MAP2K1) (PDB: 4LMN, shown in grey). **(D)** PTM-modified state of MAP2K1. Experimental position of the drug (PDB id: EUI) and Ser222 are shown in magenta and thick sticks. Drug positions modelled by RFAA shown in green, Chai-1 in orange, KarmaDock in cyan, AlphaFold3 in slate. Chai-1 model structures shown in light yellow, AlphaFold3 models in light blue. The phosphorylated residue is shown in colors corresponding to the methods by which it was modeled.

Thus, our modeling results for the test set revealed that Chai-1 and RoseTTAFold All-Atom can predict certain effects of PTMs on small molecule binding, aligning with experimental data, however some cases of Chai-1 models showed high standard deviation (Fig. 2B). AlphaFold3 showed strong accuracy in predicting ligand positions in unmodified states; however, in the majority of cases from the benchmarking dataset, PTM introduction did not alter ligand positioning. On the other hand, AlphaFold3 was the only tool which correctly captures the inactive state of insulin-like growth factor 1 receptor (Fig. 5A). Nevertheless, there are cases where AlphaFold3 predicted a significant impact of PTMs on small molecule binding (see below). In general, no method demonstrated high consistency in predicting the effects of PTMs on small molecule binding in test set, likely due to the limited availability of experimental PTM-containing structures used for training these models. However, assessing the accuracy of each method for such predictions requires a significantly larger benchmarking set, which is beyond the scope of this paper. All models obtained for the discussed test set are available for download at the DrugDomain database.

### Phosphorylation of NADPH-Cytochrome P450 Reductase, detected in two cancer types, causes significant structural disruption in the binding pocket

To generate PTM-modified protein models for the set of small molecule binding-associated PTMs identified using the DrugDomain database, we used AlphaFold3, RFAA, and Chai-1. KarmaDock was used additionally for the cases discussed in this paper. Ligand RMSD was calculated between the PTM-modified model and the experimental PDB structure or AlphaFill model, in a manner similar to that described above for the test set. LDDT-PLI score was calculated between the PTM-modified model and the experimental PDB structure. The distribution of ligand RMSD values for AlphaFold3 and RFAA is shown in Figure 6. The distribution of lDDT-PLI score is shown in Supplementary Figure 2. AlphaFold3 generates five models per run, whereas RFAA generates only one. We used all models for RMSD and lDDT-PLI score calculations. The majority of analyzed cases showed a ligand RMSD of less than 5 Å (Fig. 6) and lDDT-PLI score between 0.8 and 1.0. This can indicate two interpretations. First, both methods accurately predicted ligand position for most unmodified states of the protein. Second, the identified small molecule binding-associated PTMs do not affect ligand binding in most cases, or the selected methods detected only a small fraction of cases with this effect. Overall, number of cases with higher ligand RMSD is larger for AlphaFill models as expected (Fig. 6B, D).

**Figure 6.**
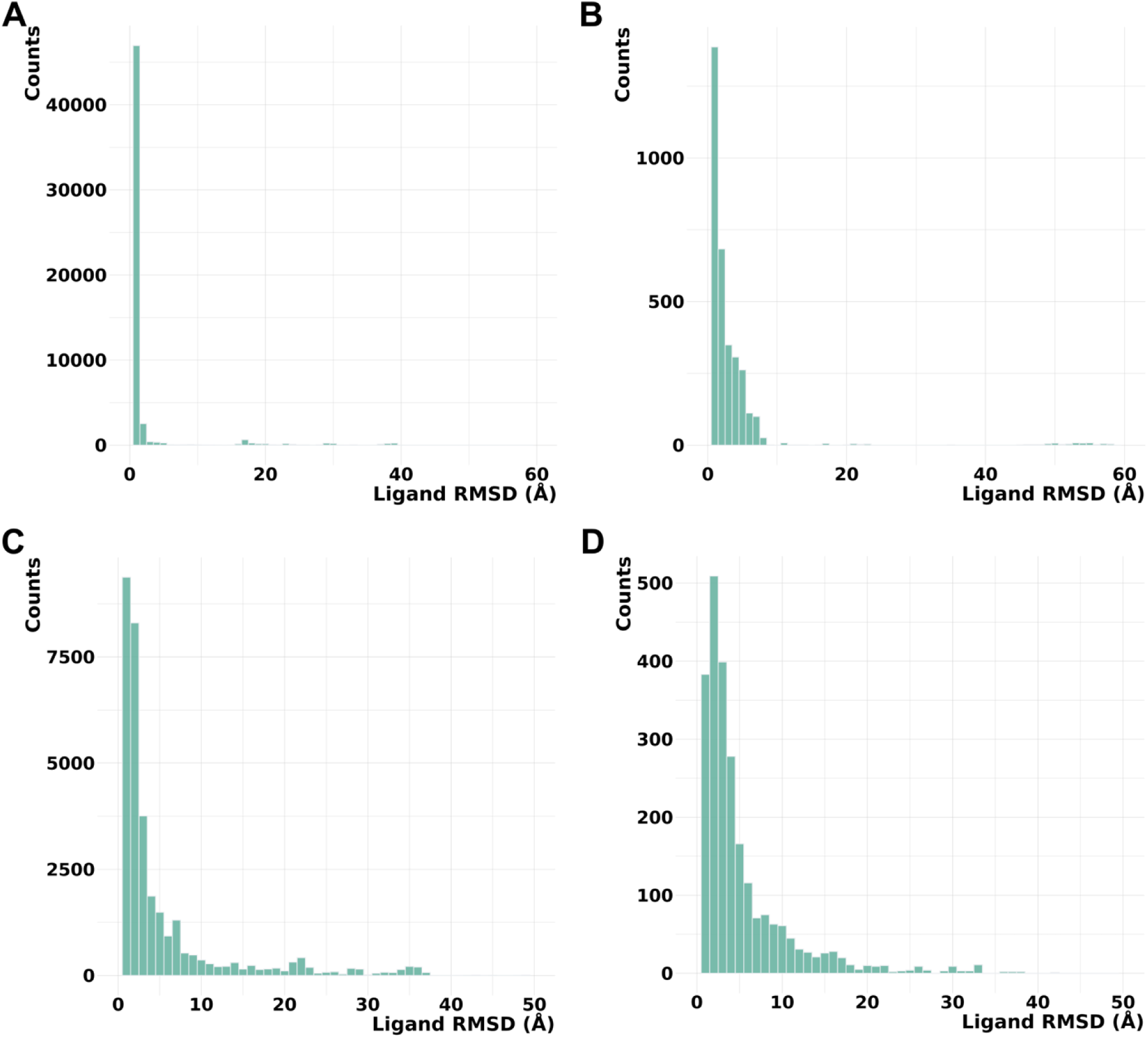
Distribution of ligand RMSD values. **(A)** AlphaFold3 models vs experimental PDB structures. **(B)** AlphaFold3 models vs AlphaFill models. **(C)** RoseTTAFold All-Atom models v experimental PDB structures. **(D)** RoseTTAFold All-Atom models vs AlphaFill models.

In many cases where the RMSD is 10–60 Å, the high RMSD value can be attributed to issue with the protein model or the specific properties of the particular small molecule. For example, RFAA did not predict the structure of the C-terminal part of the Aminoimidazole-4-carboxamide ribonucleotide transformylase (PDB: 1PL0), leading to the ligand being incorrectly positioned in the model, bound to another domain. This resulted in a ligand RMSD of 50 Å between the modeled and experimental structures (Supplementary Fig. 3). Another example is mitochondrial aldehyde dehydrogenase (PDB: 3N80) with guanidine as a ligand. In this case, guanidine is part of the solution and does not have a binding site, which resulted in high ligand RMSD values (Supplementary Fig. 4) [72]. Supplementary Figures 1 and 3 illustrate one of the major challenges that AI-based methods, including AlphaFold, have yet to overcome. While AlphaFold achieves near-experimental accuracy in predicting individual domain structures, it often struggles with accurately predicting inter-domain orientations and protein interfaces, leading to discrepancies in the overall structural arrangement [73–75]. Crucially, the ability to adopt multiple conformations is often vital for protein function, as exemplified by antibodies. However, AlphaFold struggles to adequately represent this conformational flexibility, potentially hindering accurate predictions of functional mechanisms [76]. Additionally, AlphaFold demonstrated inconsistent performance in predicting alternative protein folds, which are often essential for functional diversity. Some were rendered with low confidence, others were patently inaccurate, and a significant proportion were simply not predicted at all, indicating a substantial limitation in its ability to capture structural heterogeneity and, consequently, to accurately predict functional outcomes [77, 78]. Thus, we believe that the next crucial phase in the evolution of AI-powered protein structure prediction tools should focus on the implementation of algorithms designed to accurately represent and predict protein conformational flexibility.

Our analysis revealed several cases where many utilized methods for predicting PTM-modified protein structures suggested a significant impact on small molecule binding. For example, we discovered that phosphorylation of Tyr604 in NADPH-Cytochrome P450 Reductase most likely disrupts substrate (NADP) binding (Fig. 7). NADPH-Cytochrome P450 Reductase the electron transport from NADP to microsomal cytochromes P450 involved in steroidogenesis, xenobiotic metabolism, and monooxygenase activities like heme and squalene oxygenation [79]. The reaction of electron transfer also requires two cofactors: FAD and FMN. Thus, disfunction of this enzyme might lead to severe consequences. Several mutations in this protein have been found to be related to the development of Antley-Bixler syndrome, which is characterized by structural abnormalities of skeletal systems [80]. Phosphorylation of Tyr604 in in NADPH-Cytochrome P450 Reductase was identified during the large scale phosphoproteome analysis of two cancer cell lines: HeLa cells (cervical cancer) [81] and PC3 lung adenocarcinoma cells [82]. However, nothing else is known about the effects of this PTM. Our modelling results showed that all four utilized approaches correctly predicted NADP position for unmodified state of the protein (Fig. 7A). For PTM-modified state all approaches suggested NADP position at the two cofactor binding sites that should be occupied by FAD and FMN (Fig. 7B). AlphaFold3 and Chai-1 predicted the position of phosphorylated Tyr to be very close to that of the experimental non-modified residue, whereas RFAA’s predicted position of the PTM differs significantly (Fig. 7B, shown in green sticks). Comparison of the substrate binding pocket measurements between experimental, unmodified AlphaFold3 model and PTM-modified AlphaFold3 model showed that unmodified AlphaFold3 structure and binding mode of NADP is very close to the experimental one (Fig. 7C, D). The introduction of phosphorylated Tyr reduces the length of the binding pocket by at least 2 Å (13.8 Å in PTM-modified model vs 15.8 Å in experimental unmodified structure), which is significant enough to disrupt substrate binding (Fig. 7E). Thus, it is not surprising that this PTM has been identified in at least two types of cancer, as its potential impact on protein function could significantly influence processes critical to carcinogenesis, including metabolism, signaling, and oxidative stress.

**Figure 7.**
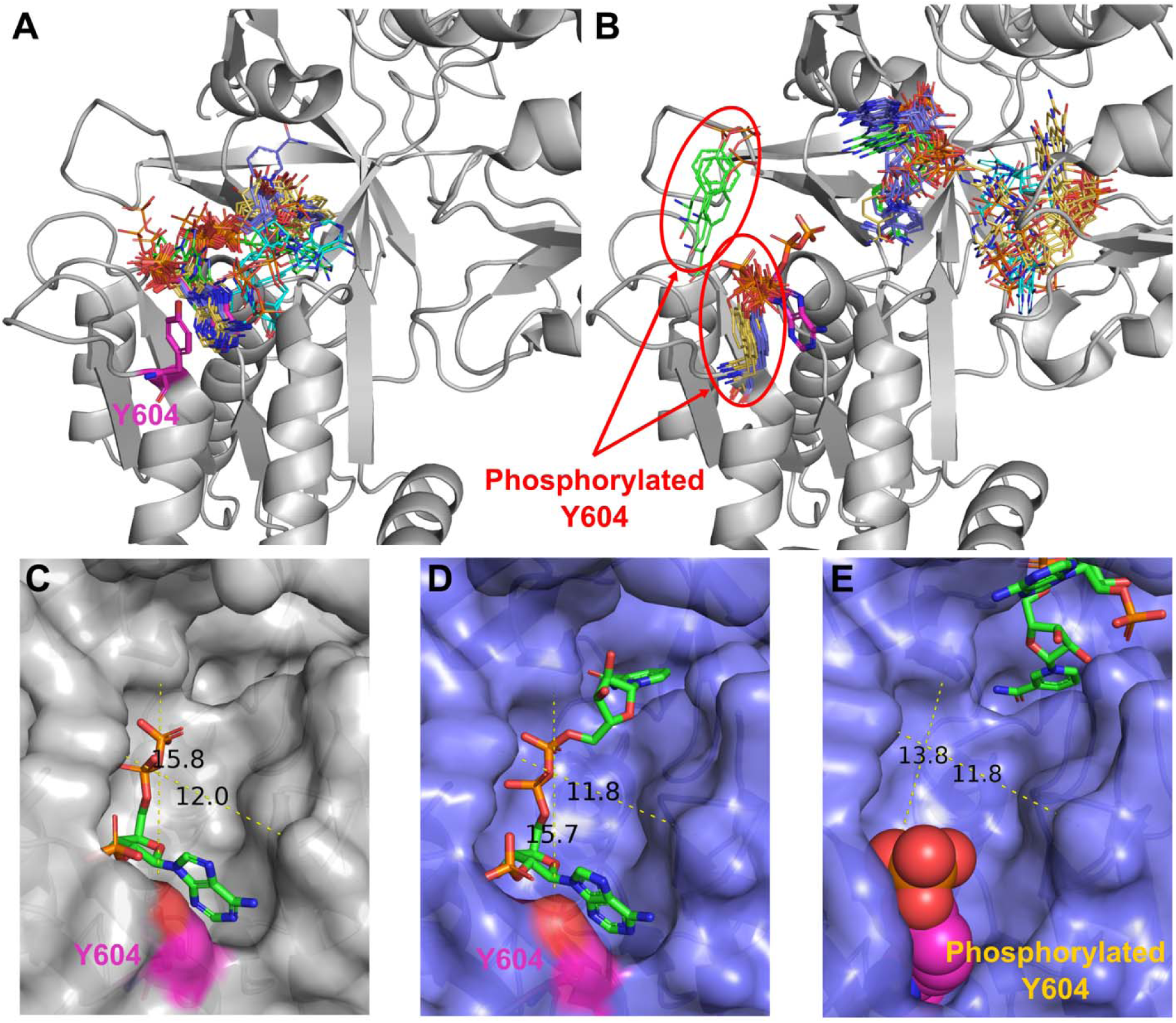
Phosphorylation of Tyr604 affects binding of NADP by NADPH-Cytochrome P450 Reductase. **(A)** Unmodified state of NADPH-Cytochrome P450 Reductase (PDB: 3QFR, shown in grey). **(B)** PTM-modified state of NADPH-Cytochrome P450 Reductase. Experimental position of the NADP and Tyr604 are shown in magenta and thick sticks. Drug positions modelled by RFAA shown in green, AlphaFold3 in purple, Chai-1 in orange, KarmaDock in cyan. The phosphorylated residue is shown in colors corresponding to the methods by which it was modeled. **(C)** Binding pocket of experimental structure of NADPH-Cytochrome P450 Reductase (PDB: 3QFR). **(D)** Binding pocket of unmodified state of NADPH-Cytochrome P450 Reductase modelled by AlphaFold3. **(E)** Binding pocket of PTM-modified state of NADPH-Cytochrome P450 Reductase modelled by AlphaFold3.

We catalogued all identified small molecule binding-associated PTMs in DrugDomain database v1.1. For each combination of protein (UniProt accession) and ligand (DrugBank ID), we provided a table of identified PTMs, if detected. This table includes information about each PTM and links to PyMOL sessions with models of modified proteins generated by AlphaFold3, RoseTTAFold All-Atom or Chai-1 (Fig. 8A). PyMOL sessions include mapped ECOD domains shown in various colors and modified residue and ligand shown in magenta (Fig. 8B, C). The complete list of identified PTMs with their corresponding ECOD domains is available for download as a plain text file from the DrugDomain website (http://prodata.swmed.edu/DrugDomain/) and GitHub (https://github.com/kirmedvedev/DrugDomain). All generated modified protein models are available for download from the DrugDomain website (http://prodata.swmed.edu/DrugDomain/download/).

**Figure 8.**
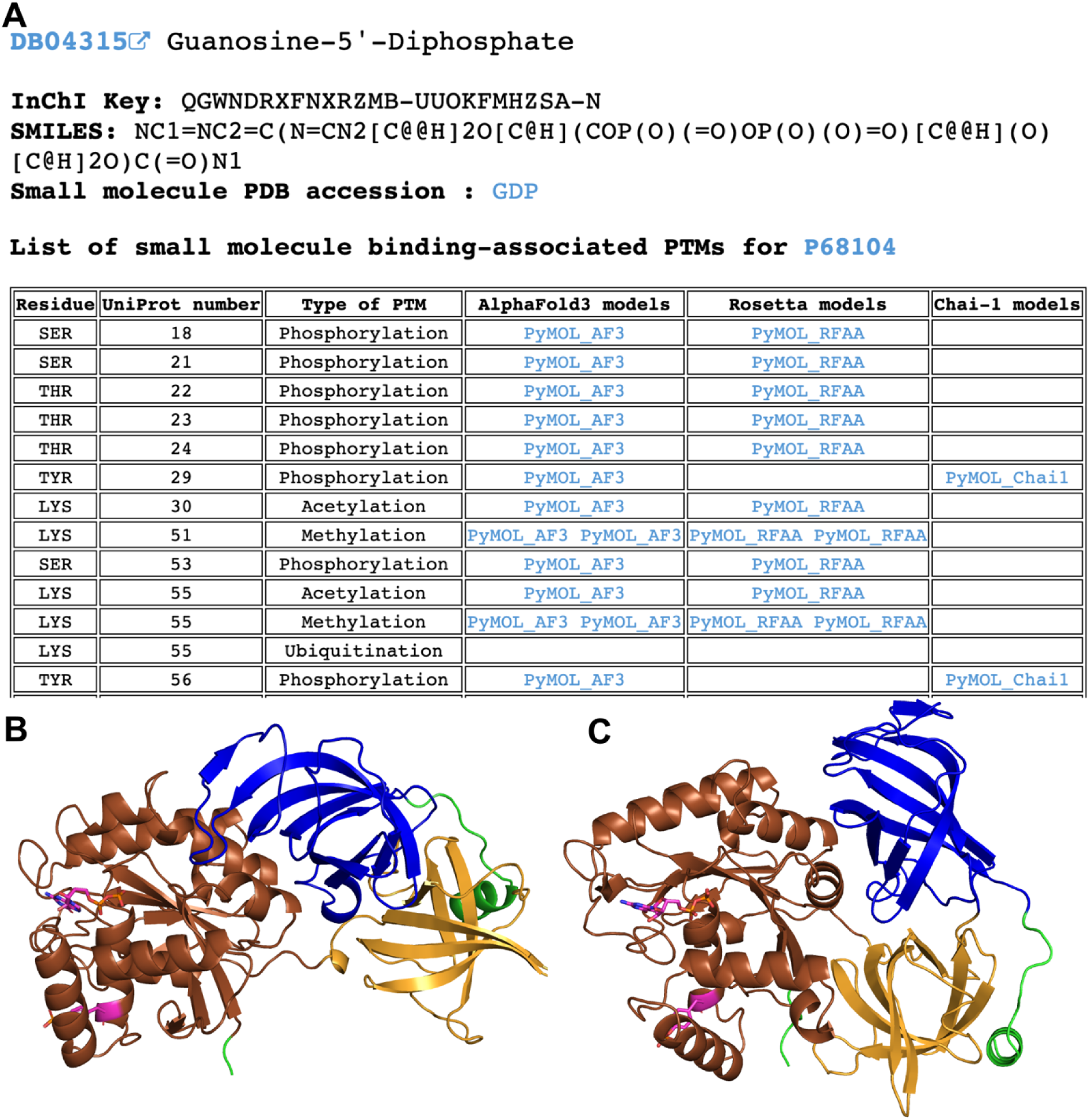
Example of the DrugDomain data webpage showing the list of small molecule binding-associated PTMs for Elongation factor 1-alpha 1 (P68104). **(A)** Table of small molecule binding-associated PTMs with links to generated models of modified structures. **(B)** AlphaFold3 model of modified structure of Elongation factor 1-alpha 1 with phosphorylated TYR29. **(C)** Chai-1 model of modified structure of Elongation factor 1-alpha 1 with phosphorylated TYR29. ECOD domains are shown in different colors.

## Conclusions

In this study, we identified post-translational modifications (PTMs) associated with small molecule binding that can influence drug binding across all human proteins listed as small molecule targets in the recently developed DrugDomain database. Mapping identified PTMs to structural domains from the ECOD database revealed that top three ECOD A-groups with the largest number of small molecule binding-associated PTMs across experimental PDB structures include a/b three-layered sandwiches (Rossmann fold), a+b complex topology (kinases), and a+b two layers (heat shock proteins). Evaluation of AI-based protein structure prediction approaches (AlphaFold3, RoseTTAFold All-Atom, Chai-1, KarmaDock) in the context of PTM structural effects revealed that Chai-1 and RoseTTAFold All-Atom can predict certain effects of PTMs on small molecule binding, consistent with experimental data. AlphaFold3 demonstrated strong accuracy in predicting ligand positions in unmodified states; however, in most cases from the benchmarking dataset, the introduction of PTMs did not affect ligand positioning. Using advanced AI-based protein structure prediction methods (AlphaFold3, RoseTTAFold All-Atom, Chai-1), we created 14,178 models of PTM-modified human proteins with docked small molecules. This data revealed cases of significant impact of PTMs on small molecule binding. For example, we discovered that phosphorylation of NADPH-Cytochrome P450 Reductase, observed in cervical and lung cancer, leads to substantial structural disruption in the binding pocket, potentially hindering protein function. We reported all identified small molecule binding-associated PTMs and all generated PTM-modified models along with test set models in DrugDomain database v1.1 (http://prodata.swmed.edu/DrugDomain/) and GitHub (https://github.com/kirmedvedev/DrugDomain). We believe this resource, to our knowledge the first to provide structural context for small molecule binding-associated PTMs mapped to structural domains on a large scale, could serve as a valuable tool for studying the evolutionary and structural aspects of PTMs.

## Acknowledgments

The authors acknowledge the Texas Advanced Computing Center (TACC) at The University of Texas at Austin (http://www.tacc.utexas.edu) for providing computational resources that have contributed to the research results reported within this paper. This research was carried out in part using the computational resources provided by the BioHPC computing facility of the Lyda Hill Department of Bioinformatics, UT Southwestern Medical Center, TX (https://portal.biohpc.swmed.edu).

## CRediT Author Contribution

**Kirill E. Medvedev:** Conceptualization, Methodology, Software, Validation, Formal analysis, Investigation, Data Curation, Visualization, Writing - Original Draft, Writing - Review & Editing, Project administration. **R. Dustin Schaeffer**: Writing - Review & Editing, Funding acquisition. **Nick V. Grishin:** Conceptualization, Resources, Funding acquisition, Writing - Review & Editing.

