## Supplementary Figures 1-4 for "Leveraging AI to Explore Structural Contexts of Post-Translational Modifications in Drug Binding"

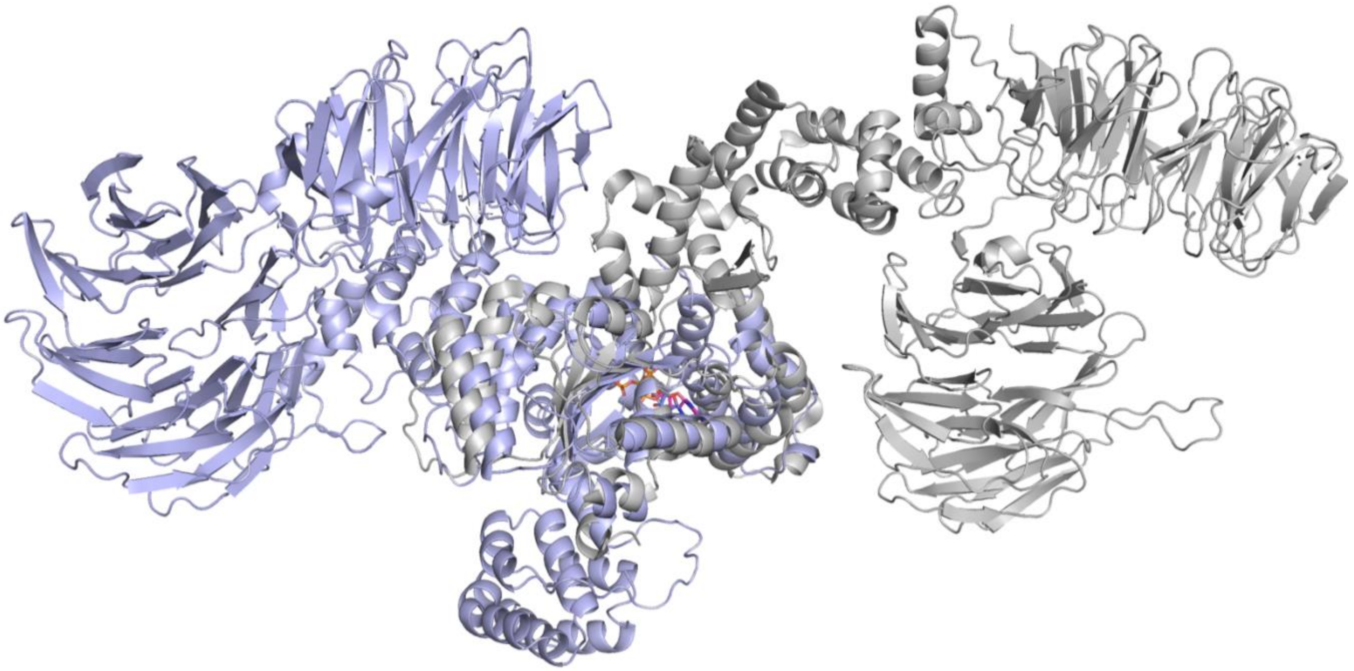

**Supplementary Figure 1. Example of mismatch of domains orientation between model and experimental structure.** Structure of apoptotic protease-activating factor 1 (PDB: 3J2T, shown in grey). AlphaFold3 model shown in light blue.

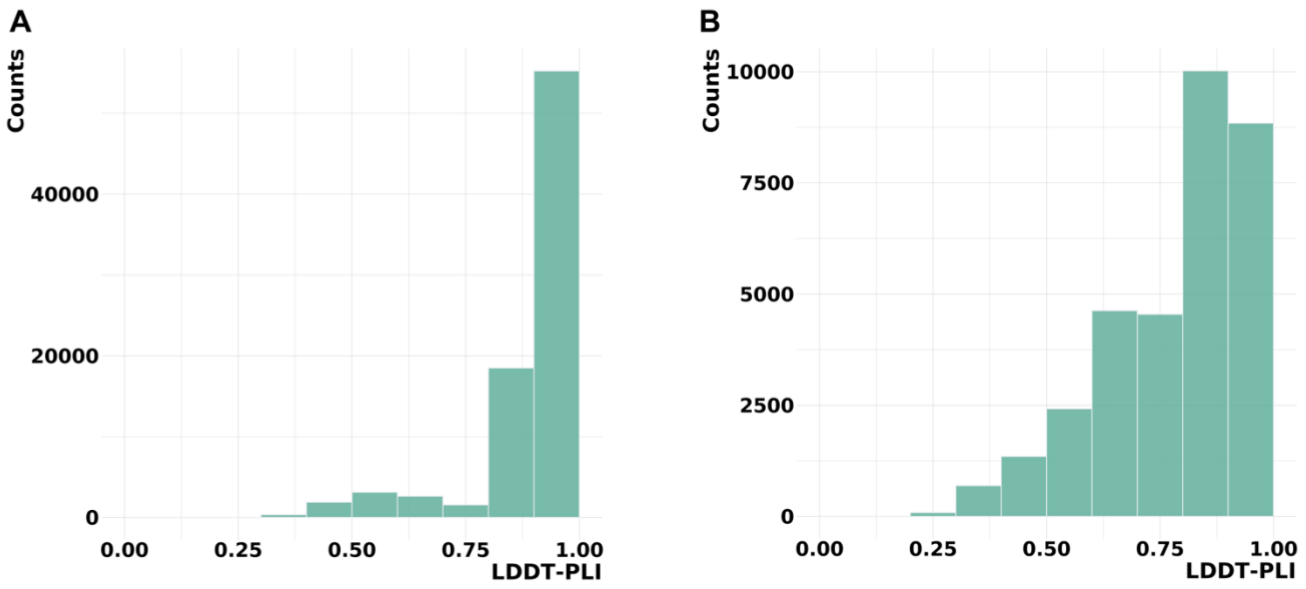


**Supplementary Figure 2. LDDT-PLI scores distribution. (A)** AlphaFold3 models vs experimental PDB structures. **(B)** RoseTTAFold All-Atom models vs experimental PDB structures.

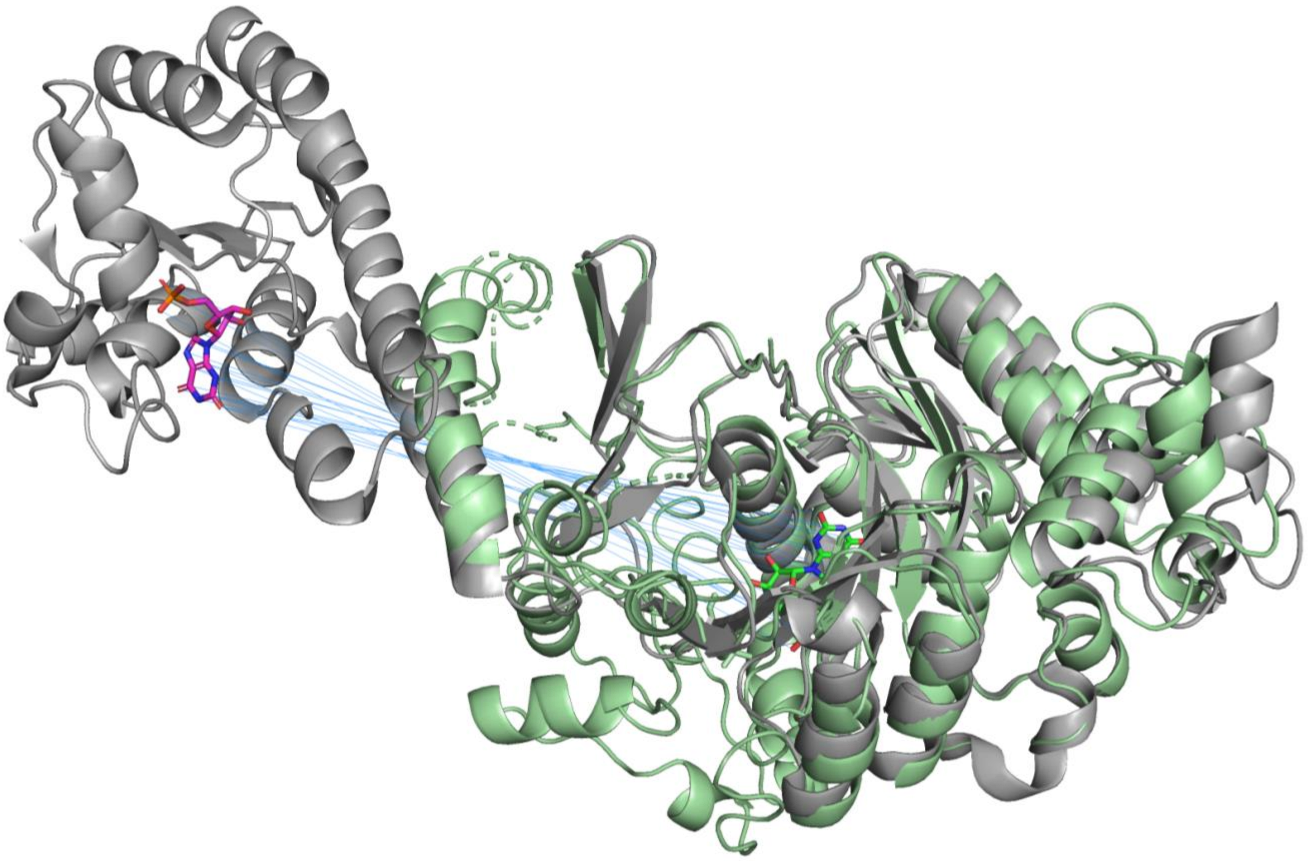

**Supplementary Figure 3. Structure of Aminoimidazole-4-carboxamide ribonucleotide transformylase (PDB: 1PL0, shown in grey).** RFAA model shown in light green. Ligand from experimental structure shown in magenta. Alignment lines between experimental and modelled ligands shown in blue.

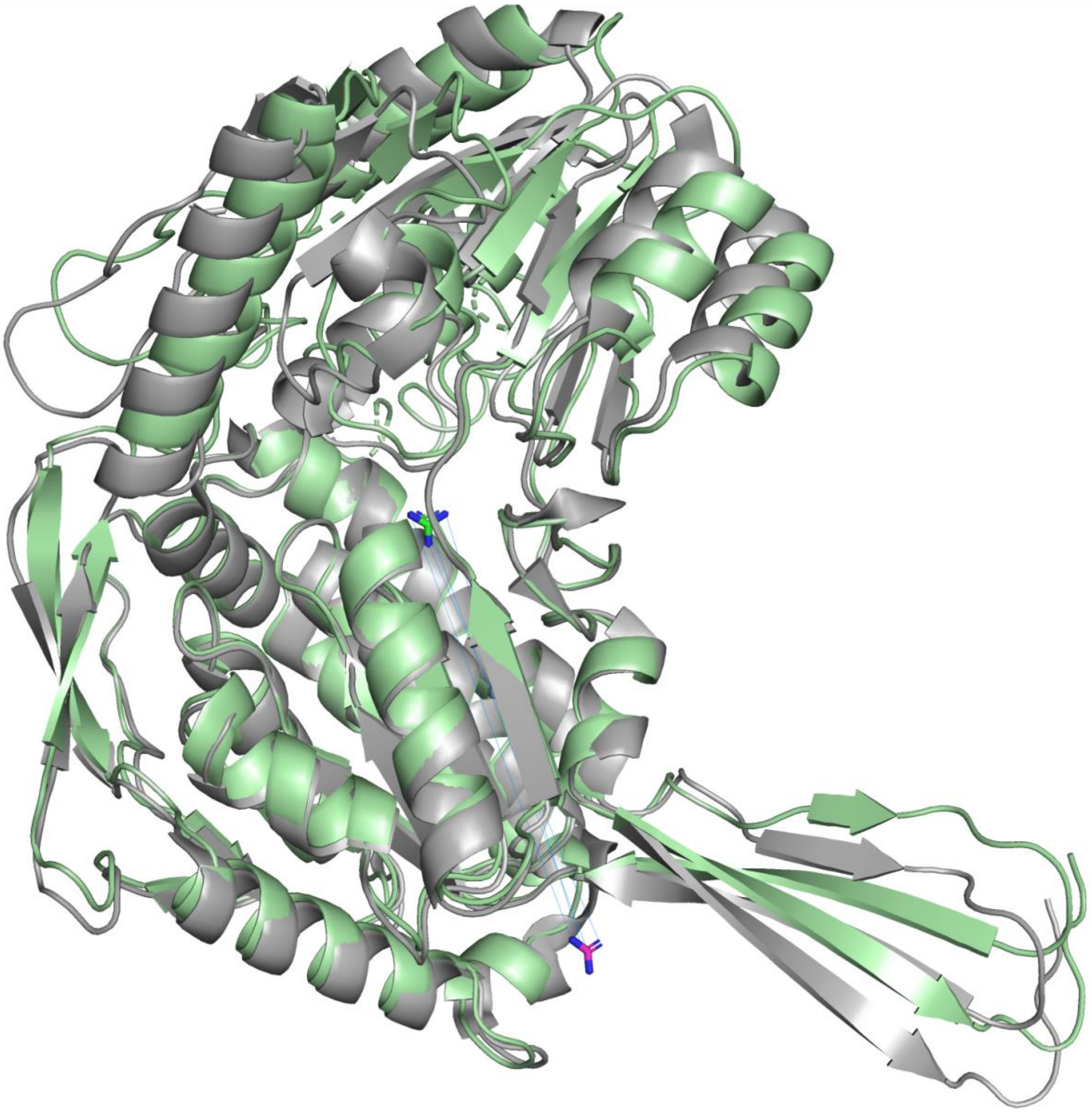

**Supplementary Figure 4. Structure of mitochondrial aldehyde dehydrogenase (PDB: 3N80, shown in grey).** RFAA model shown in light green. Ligand from experimental structure shown in magenta. Alignment lines between experimental and modelled ligands shown in blue.
